# Drought, thermal response and climate-patterning in cuticular conductance of the widespread C4 grass, *Themeda triandra*

**DOI:** 10.1101/2025.11.02.686155

**Authors:** Anu Middha, Vinod Jacob, Chris J. Blackman, Brendan Choat, Ian J. Wright

## Abstract

- *Background and Aims*: Grasslands underpin global biodiversity, carbon storage, and ecosystem resilience, yet are increasingly threatened by rising temperatures and water scarcity. Understanding how key physiological traits respond to drought and heat is essential for ensuring grassland function under future conditions. Here, we investigated intraspecific variation in leaf cuticular traits—specifically minimum cuticular conductance after stomatal closure (g_min_) and its response to temperature—in six Australian accessions of *Themeda triandra* spanning a wide climatic gradient. We asked whether g_min_ and its response to drought reflect climate-of-origin and whether the cuticle shows a thermal threshold (Tp) beyond which conductance rises sharply.
- *Methods*: Plants from six accessions were grown under well-watered (control) glasshouse conditions and then exposed to drought. We measured g_min_ in both control and drought treatments and assessed the response of gmin to increasing temperature (30−55 °C) using fresh fully hydrated leaves (control). Climatic data for each site of origin were used to explore trait−environment relationships.
- *Key results:* Under well-watered conditions, g_min_ showed no link to climate-of-origin. Under drought, however, g_min_ displayed clearer climate-linked patterns: accessions from cooler, wetter regions had lower values, contrary to expectations. Drought responses varied strongly among accessions, ranging from marked reductions to significant increases in g_min._ With increasing temperature, g_min_ declined gradually and no accession exhibited a distinct phase transition, indicating a thermally stable cuticle.
- *Conclusions*: *Themeda triandra* shows considerable intraspecific diversity in g_min_ and its response to drought. The clearer alignment between climate of origin and g_min_ measured under drought conditions suggests drought stress is an important filter for g_min_ expression. These findings provide a physiological basis for identifying genotypes with enhanced resilience for use in grassland conservation and restoration under a warming, drying climate.

## Introduction

Grasslands occupy 31% of earth’s land area (Petri *et al*., 2010) and are essential for preserving biodiversity, sequestering carbon and maintaining ecosystem balance (Blair *et al*., 2014). However, they are highly vulnerable to climate change, particularly rising temperatures and altered rainfall patterns (Sala *et al*., 2000, Teuling, 2018). These shifts threaten grassland resilience by reducing species diversity, productivity, and soil integrity (Overpeck, 2013, Edenhofer *et al*., 2014), underscoring the need to understand their impacts for effective conservation and management.

Under well-watered daytime conditions, stomata remain open to facilitate gas exchange for photosynthesis while simultaneously losing water via transpiration. During drought, plants reduce water loss primarily by closing their stomata. However, even after full stomatal closure, plants continue to lose water through the leaf epidermis (“residual transpiration”; Billon *et al*. (2020)). This residual transpiration can be significant, particularly under prolonged drought or high temperatures, and is primarily attributed to the permeability of the leaf cuticle, although leaky stomata may also contribute (Muchow & Sinclair, 1989, Machado *et al*., 2021). The rate of this residual water loss from the leaf surface when stomata are fully closed is often referred to as “minimum cuticular conductance” or “g_min_”. g_min_ is increasingly recognized as being ecologically important, indexing the effectiveness of the cuticle for slowing water loss during drought (Schuster *et al*., 2016). In dryland and savanna species, for example, small differences in g_min_ can significantly affect drought survival (Burghardt & Riederer, 2003). Recent work has highlighted that g_min_ is not constant but dynamic throughout dehydration (Burlett *et al*., 2025). This variability in g_min_ can potentially alter predictions of time to hydraulic failure (Martin□StPaul *et al*., 2017), thus highlighting the need to understand the response of g_min_ to drought stress, especially against the background of projected increasingly prolonged and severe droughts.

At moderate temperatures (e.g. 20−25 □), g_min_ of healthy intact leaves is relatively low, typically averaging around 10% of the transpiration through fully open stomata, depending on the species (Holmgren *et al*., 1965, Boyer *et al*., 1997). This is because cuticular waxes serve as an efficient barrier against water movement (Boyer *et al*., 1997, Burghardt & Riederer, 2003, Riederer, 2006). However, g_min_ may vary with ambient temperature, displaying non-linear dynamics such that it remains relatively stable across these moderate temperatures, then increases sharply beyond some threshold temperature (Schreiber & Schönherr, 1990). This threshold temperature is commonly known as the “phase transition temperature” or (T_p_). At T_p_, the structural integrity of the cuticle begins to shift due to thermal disruption of wax and polymer organization, resulting in a dramatic increase in g_min_ and hence, higher water loss (Schreiber & Schönherr, 1990). T_p_ can vary among species and populations and is considered a marker of thermal stability of the cuticle (Schuster *et al*., 2016, Bueno *et al*., 2019, Billon *et al*., 2020). For example, Schuster *et al*. (2016) reported that the desert shrub *Rhazya stricta* had a markedly high T_p_, suggesting an adaptation to hot, arid conditions. The temperature response of g_min_ is further influenced by the chemical composition of cuticular waxes, as compounds such as alkanes, fatty alcohols, aldehydes and cyclic triterpenoids differ in their melting behaviour and structural stability under heat. Busta *et al*. (2021) showed that triterpenoid-rich sorghum leaves possess a stronger, more heat-resistant cuticular barrier than maize leaves which lacked triterpenoids. In the context of climate change and heatwaves, understanding T_p_ is critical because it determines the threshold temperature at which the primary barrier to plant water loss is compromised during drought. The extent to which these changes are reversible is also unknown. For instance, Schönherr *et al*. (1979) observed that heating cuticular membranes above their transition temperature irreversibly altered the distribution of soluble cuticular lipids, leading to increased water permeability.

Substantial interspecific variation in g_min_ has been documented; e.g. Duursma *et al*. (2019) compiled data for 221 species from 40 original studies, reporting values from ∼ 0.1 to 50 mmol m^−2^ s^−1^. By taxonomic order, Poales on average exhibited the highest values, and conifers (Pinales, Araucariales) the lowest, but there was wide variation within any given group. There is evidence showing that g_min_ variation reflects differences in the chemical composition of cuticles; e.g. the richness of cuticular wax components (such as long-chain alkanes) is known to reduce cuticle permeability (Huang *et al*., 2017, Cheng *et al*., 2019).

Climate-related *intraspecific* variation in g_min_ and cuticle traits has also been observed. For example, some studies have reported g_min_ as lower in plants from arid compared to mesic environments, presumably to minimize residual water loss during drought; this was the case for one of two *Hakea* (Proteaceae) species studied by López *et al*. (2021). Furthermore, Smith *et al*. (2006) compared *Digitaria californica* populations from Texas (arid) and Arizona (less arid), as well as the invasive species *Eragrostis lehmanniana*, in a common garden setting. They found that the arid-zone population and *E. lehmanniana* both exhibited significantly lower g_min_ values. In *Nicotiana benthamiana*, two accessions exhibited contrasting drought strategies: plants from arid central Australia relied on rapid growth and early flowering to escape drought, with minimal changes in cuticle properties or g_min_ under experimental drought (Asadyar *et al*., 2024). In contrast, plants from tropical northern Australia reduced g_min_ in response to water limitation by altering wax composition, specifically increased very-long-chain alkanes, to minimize water loss under drought conditions (Asadyar *et al*., 2024). Other studies have also shown acclimation of g_min_ to drought and elevated temperature conditions (Duursma *et al*., 2019). However, other studies have shown a lack of response in g_min_ to changing growth conditions (Bueno *et al*., 2020).

Most research on cuticle conductance has focused on woody species and a limited subset of herbaceous plants, primarily high-value annual crops such as wheat and sorghum (Muchow & Sinclair, 1989, Araus *et al*., 1991, Qariani *et al*., 2000, Lin *et al*., 2020). In contrast, native and perennial grass species—despite their ecological significance—remain understudied. As a result, our understanding of how perennial grasses regulate cuticular water loss in response to drought remains limited, highlighting the need for broader investigation across diverse grass lineages.

This study focuses on *Themeda triandra*, a perennial C4 tussock grass endemic to Africa, Australia and Asia (Snyman *et al*., 2013). It has great ecological significance due to its broad distribution across various ecosystems and geological substrates. It is a valuable source of food for both native and introduced herbivores, and plays a crucial role in supporting rural livelihoods, wildlife and livestock production (Snyman *et al*., 2013). *T. triandra* is widely distributed across Australia and is believed to have been the primary dominant species in many Australian grasslands prior to European settlement (Mitchell & Miller, 1990). It grows in habitats as diverse as the hot, semiarid interior, the wet tropics, and cool temperate sub-alpine regions (Mitchell & Miller, 1990). This makes it an ideal model for exploring physiological responses to climate-induced stresses in grasses. A reduction in the occurrence of *T. triandra* within grasslands is often associated with a drop in grazing potential, a loss of biodiversity, reduced vegetation cover and a decline in overall health of the ecosystem (Snyman *et al*., 2013). Despite its huge ecological and economic significance, to this point there has been no attempt to study its cuticle properties.

In this study, we measured g_min_ in plants under well-watered and droughted conditions and explored its response to increasing temperature across six distinct accessions of *T. triandra* originating from diverse regions across Australia, spanning a wide range of temperature and rainfall regimes. We had two key aims. First, we sought to elucidate the relationships between climate conditions at the seed source (“climate-at-origin”) and two aspects of cuticle permeability: 1) g_min_ and 2) its temperature response (T_p_). Second, we sought to characterise the different accessions in terms of their variation in g_min_ and T_p_ under control and water-limiting conditions.

We tested the following hypotheses:

1. *T. triandra* accessions from hotter and drier environments would exhibit generally lower g_min_, reflecting selection for reduced cuticular water loss under high evaporative demand and limited water availability.
2. Cuticle conductance would show a bi-phasic response to increasing temperature, remaining relatively steady at moderate temperatures, then rising sharply once temperatures exceeded a threshold, T_p_, which itself would vary with climate-at-origin. Specifically, we expected plants from hotter environments to have higher T_p_; i.e., to maintain cuticular stability up to higher temperatures.
3. Experimental drought would reduce g_min_ in all accessions. This expectation assumes that the lower gmin would reduce plant water loss when savings are needed most. We also expected the proportional change in g_min_ (from control to drought conditions) to be greater in accessions originating from arid or high-temperature regions. This is based on the assumption that plants from such environments have evolved greater plasticity or regulation of cuticle properties to limit water loss during droughts (Bueno *et al*., 2019).

## Materials and methods

### Plant material and growth conditions

Six accessions of *T. triandra* (Dalby, QLD; Forbes, NSW; Mt. Fox National Park, Queensland, Pannawonica, Western Australia; Rainbow Valley, Northern Territory; and Virginia Gardens, South Australia) (Fig. 1 (a)) were grown from seeds sourced from the Australian Pastures Genebank, Native Seeds Pty Ltd and Nindethana Seed Service Pty Ltd (Albany, Western Australia). The selected accessions represent a range of climate-at-origin conditions (climate data from CHELSA (Karger *et al*., 2017); Table S1), with mean summer temperatures ranging from 21.0 □ to 32.0 □, summer rainfall from 75.1 to 435.9 mm, annual temperatures (AT) from 16.2 □ and 22.1 □ and annual precipitation (AP) from 281 to 712 mm. Plants were propagated from seeds, germinated in seed trays, and transplanted into 250 mm (∼8 L) pots filled with sandy-loam soil (volumetric water holding capacity of 15−20%, pH ≈ 5.6) collected from the Pastures and Climate Extremes (PACE) field facility at Western Sydney University (Churchill *et al*., 2022), supplemented with slow-release native fertiliser (Osmocote^®^ Native; 20 g pot^−1^)

**Figure 1.**
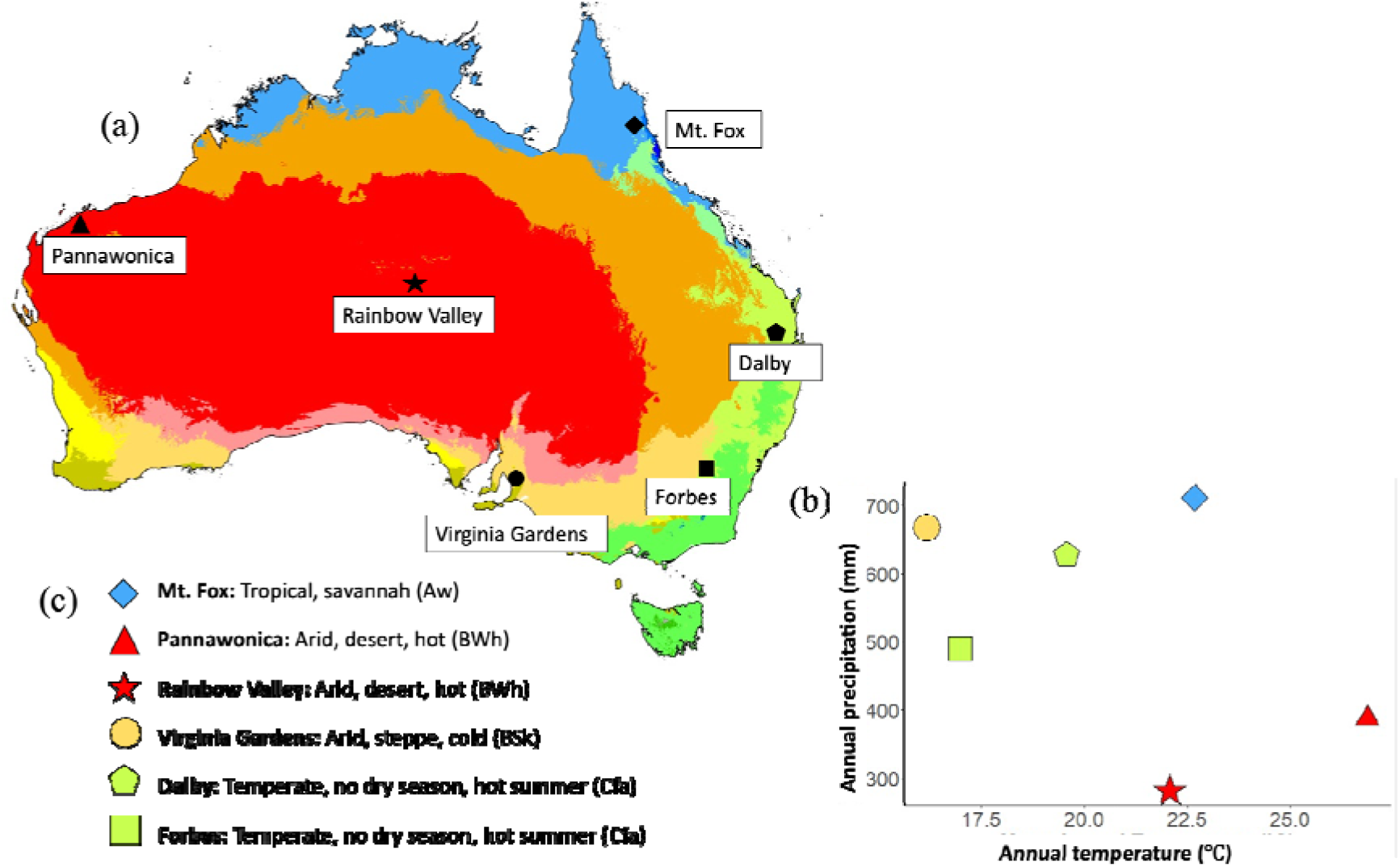
(a) Köppen-Geiger climate classification map of Australia adapted from Peel *et al*. (2007) showing the distribution of various climate zones. The locations of six *Themeda triandra* accessions included in this study are marked on the map. (b) Scatterplot illustrating the annual temperature (AT) and annual precipitation (AP) of the selected accessions, highlighting the climatic differences across their native habitats. (c) Köppen-Geiger climate classifications represented by distinct colours corresponding to specific climate types based on temperature and precipitation patterns. (The Köppen-Geiger classification system utilizes a combination of letters and colours to denote different climate types. For instance, ‘BWh’ in red denotes a hot desert climate with high temperatures and minimal precipitation. These classifications are visually represented through distinct colour codes to facilitate easy identification on the map)

### Water deficit treatment

Each accession was represented by four replicates, all maintained under well-watered conditions for two months. Plants were irrigated to full field capacity using a drip system. Toward the end of this period, all four replicates from each accession were sampled for g_min_ and Tp measurements under well-watered conditions. The plants were then subjected to an acute water-deficit treatment by withholding irrigation until the leaves were visibly wilted (3 days for all plants) to simulate a rapid decline in soil moisture. Soil water content was monitored using a soil moisture probe, and leaf water potential was measured with a pressure chamber to assess plant water status (Figs S1 and S2). At the end of the drying period, the plants were re-sampled for determination of gmin under water-deficit-treated conditions.

### Leaf drying methods used for g_min_ determination

Initially, g_min_ measurements were conducted using the benchtop drying method (Sack & Scoffoni, 2010) at a single temperature (30 □), which was well-suited for assessing the steady state cuticular conductance under controlled conditions. However, as the study progressed, the need to measure g_min_ across multiple temperatures became evident, requiring a more efficient and automated approach. The benchtop method, though reliable, was time-and labour-intensive and limited in its throughput. To overcome these limitations and facilitate rapid measurements across a range of temperatures, we incorporated a “DroughtBox” (Billon *et al*., 2020) into our workflow, which provided an automated, high-throughput system for assessing g_min_ under precisely controlled environmental conditions.

### Description of the DroughtBox

The DroughtBox, originally described by Billon *et al*. (2020) consists of a 70 cm cubic box with two compartments for semi-automatically determining g_min_ under controlled environmental conditions, specifically temperature and humidity. The system includes strain gauges connected to a data acquisition board (for measuring the mass loss from leaves as they dehydrate), heating resistors for temperature control, and electronically controlled valves for adjusting relative humidity. The original design also incorporates a Raspberry Pi for overall control and monitoring.

In this study, we used a modified version of the DroughtBox to measure g_min_ and T_p_. Our version consisted of a 60 cm × 140 cm × 70 cm cuboidal box made of 40 mm thick polystyrene boards with a front door and two compartments (Fig. S4). The upper compartment housed 8 strain gauges (Phidgets single point load cell 780g), fixed in an aluminium frame and directly connected to a Campbell Scientific CR300 datalogger for data acquisition. Leaves for g_min_ measurements were placed in the main lower compartment and attached to the strain gauges through holes in the roof board. An air temperature and relative humidity sensor (Vaisala HMP155) was suspended in the lower compartment to monitor environmental conditions, along with a fine-wire type T thermocouple for rapid response temperature control. Temperature regulation was achieved using two PTC heaters (Joyzan, 220 V, 300 W) mounted in galvanized steel conduits, with a 50 mm 12 V fan to push heated air into the box. A thermostat switch (MULTICOMP T23A150BSR2-15) was installed for safety, shutting off the heaters if they overheat. A solid-state relay (Sparkfun electronics, 40A, 3−32 V DC) was used to allow logger to regulate the heaters via pulse width modulation. Relative humidity in the lower compartment was adjusted by a water-filled plastic container with ultrasonic mist maker (Vaguelly mist maker, 350 mL, 24 V). A 12 V pump (DANXQ, 12 V DC) circulated air through the mist container to introduce fine mist into the box when needed. All environmental variables were monitored and controlled using a Campbell Scientific CR3000 datalogger with program written and compiled in Visual Basic.

### Performance testing of the DroughtBox system

The ability of the DroughtBox to achieve and maintain stable temperature and humidity conditions was assessed. To do this, the system was programmed to run at each target temperature and relative humidity required for our experiment over 24-hour periods, with temperature, relative humidity, and VPD recorded every 10 seconds.

### Cuticle conductance measurement (benchtop drying)

To assess leaf water loss, 4−5 healthy tillers with fully developed, mature leaves were collected in the morning on each sampling day. The cut ends of the tillers were immediately wrapped in damp paper towels, placed in plastic zip lock bags, and transported to the lab. To relieve xylem tension, the tiller ends were recut under water and allowed to rehydrate in falcon tubes for about 30 minutes. The cut ends were then sealed with parafilm. The tillers were weighed using a balance to record their fully hydrated weight. The leaves were then dehydrated in open plastic trays placed in a controlled growth cabinet set at 30 °C and 60% relative humidity, in the dark, for 2−3 hours, with leaf weight (w) measured every 20 minutes until the rate of water loss stabilized. We considered changes in w to represent the number of water molecules (n_w_, mmol) transpiring from the branches. Based on this assumption, transpiration rate, E (mmol m^−2^ s^−1^) was calculated as

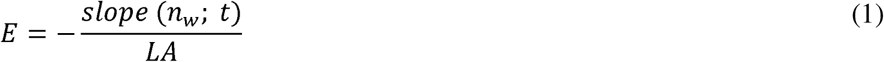

where, t is the time in seconds, and LA represents two-sided leaf area. LA was estimated using the dry leaf mass of each tiller set and empirical relationships between leaf dry mass and fresh leaf area (i.e., LMA). Subsequently, g_min_ was determined from the linear segment of the water loss curve, that is, after stomatal closure, as following

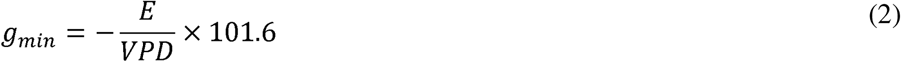

where, E is transpiration in mmol m^−2^ s^−1^, VPD refers to the vapour pressure deficit of the air in kPa, and 101.6 denotes the atmospheric pressure within the cabinet (described below) in kPa.

At the end of the experiment, the leaves were oven-dried at 70°C for 72 hours, and the final dry weight was recorded.

The percentage change in g_min_ in response to drought (%Δg_min_) was determined using the g_min_ data from control and drought-treated plants. It was calculated using the formula

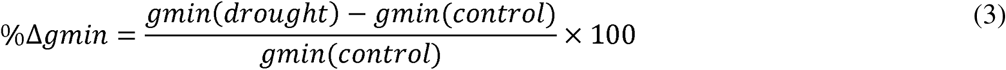

### g_min_ measurement using the DroughtBox

The tillers were harvested and prepared for drying in a manner consistent with benchtop drying procedure. Multiple leaf-drying runs were conducted in the DroughtBox at target temperatures of 30, 35, 40, 45, 50 and 55 □ (using a distinct cohort of samples at each temperature), with corresponding increases in relative humidity (RH) to maintain a constant absolute humidity of 18 g(H_2_O)m^−3^. This standardization ensured consistent conditions during g_min_ measurements. The mass data from the DroughtBox were similarly used to calculate g_min_ from the linear portion of the mass loss curve, as was done following benchtop drying procedure. After the DroughtBox measurements were complete, samples were oven-dried at 70 □ for 72 hours to determine their dry mass.

### Leaf mass per area (LMA)

LMA was calculated as the ratio of leaf dry mass to leaf area. For each replicate plant, fully expanded leaves were harvested and scanned using a LICOR leaf area meter (model LI-3100C, LI-COR Biosciences, USA) to determine the total leaf area. The scanned leaves were then oven-dried at 70 °C for 72 hours to reach dry mass. LMA (g m^−2^) was then calculated by dividing dry mass by leaf area.

### Statistical analyses

Differences in midday leaf water potentials were analysed using two-way ANOVA. To assess treatment effects within each accession, assumptions of normality (Shapiro-Wilk test) and homogeneity of variance (F-test, Levene’s test) were evaluated. Based on these. Appropriate statistical tests (paired t-test, Welch’s t-test, or Wilcoxon signed-rank test) were used. The relationship between leaf water potential and the direction and magnitude of the g_min_ response across individuals was analysed using ordinary least squares linear regression.

The g_min_ data, including those obtained from both benchtop drying and from multiple temperature runs in the DroughtBox, were visually examined for normality before analysis using histograms and density plots, and log-transformed wherever the data were right-skewed. Significant differences in g_min_ values between accessions and treatments were assessed using a two-way analysis of variance (ANOVA), followed by Tukey’s Honestly Significant Difference (HSD) test and paired t-test for pairwise comparisons. The relationships of g_min_ and %Δg_min_ with climate variables were evaluated through multiple linear regression models using mixed effects models with accession treated as a random effect. By including both temperature and rainfall in these models, we were able to identify the rainfall-independent effect of temperature and the temperature-independent effect of rainfall. Climate variables were treated both on a yearly basis (annual temperature, AT; annual precipitation, AP), and on a seasonal basis (mean summer temperature; summer rainfall).

To determine T_p_, we used segmented regression applied to Arrhenius-transformed data. Specifically, g_min_ values were natural log-transformed (ln g_min_) and plotted against the inverse of absolute temperature (1/T, in Kelvin). For each replicate, a segmented linear regression model was fitted using the segmented package in R to estimate a potential breakpoint along the temperature axis, corresponding to a putative T_p_. A significance threshold of 0.05 was applied for all statistical tests.

All analyses were conducted using R v.4.4.0 (R, **Core Team** 2025).

## Results

### DroughtBox performance under different temperature settings

When programmed to maintain constant temperatures of 30, 35, 40, 45, and 50 °C—with corresponding increases in RH to keep absolute humidity consistent—the DroughtBox successfully maintained stable temperature, relative humidity, and VPD throughout each 24-hour run (Fig. S5).

### Analysis of drought exposure and effect on g_min_

Although soil moisture declined at different rates among accessions and replicates (Fig. S1), indicating that plants experienced drought progression at slightly different rates, all droughted plants exhibited visible signs of leaf wilt and crossed their accession-specific TLP (Jacob *et al*, unpubl. res.) (Fig. S2), confirming that each plant experienced physiological drought. In all accessions, droughted plants exhibited a decline in leaf water potential relative to control conditions (ANOVA: treatment, p < 0.001, accession x treatment, p < 0.05; pairwise differences: all p < 0.05) and all crossed their respective TLP indicating loss of turgor. We found no significant relationship between percentage change in g_min_ from control to drought (%Δg_min_) and leaf water potentials (p = 0.642, R^2^ = 0.01).

### Drought-induced changes in cuticular conductance across accessions

Pooling data across the six accessions, g_min_ did not differ between control and droughted plants (Fig. 2). However, substantial variation among accessions was evident, both in mean g_min_ values and in their response of g_min_ to drought (Fig. 3). g_min_ of control plants varied among accessions (ANOVA, p < 0.001), being significantly higher in plants from Pannawonica and Mt. Fox than those from Dalby (Tukey HSD, both p < 0.001), Forbes (Tukey HSD, p = 0.003 and 0.015, respectively), and Virginia Gardens (Tukey HSD, p = 0.004 and 0.019, respectively).

**Figure 2.**
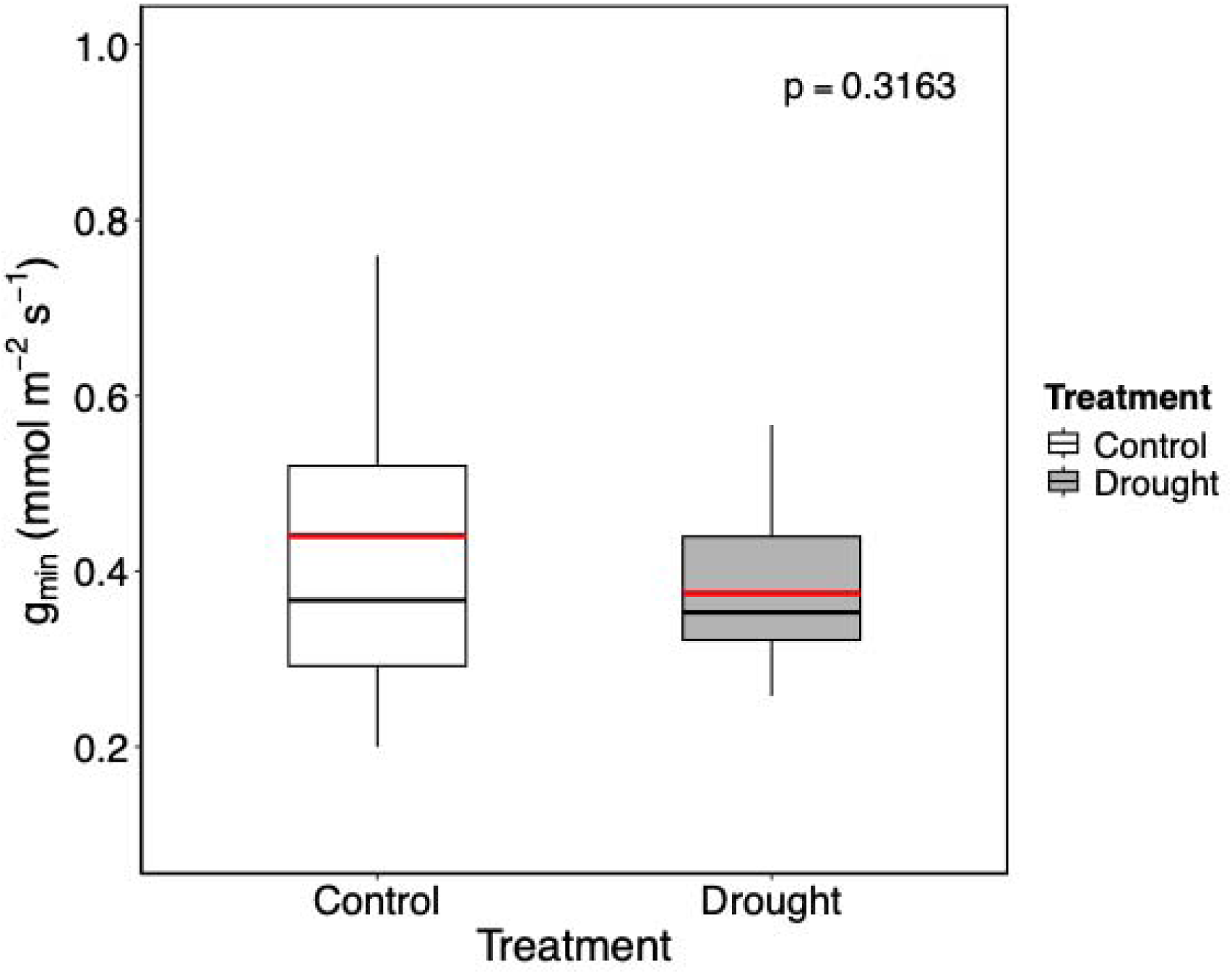
Box plot showing the distribution of g_min_ under control and drought treatments. The boxes represent the interquartile range (IQR), the black horizontal line within each box represents the median, and whiskers extend to 1.5 × IQR (n = 24). Red line inside the boxes represent mean. The p-value (0.3163; paired t-test) reflects the statistical comparison between treatments.

**Figure 3.**
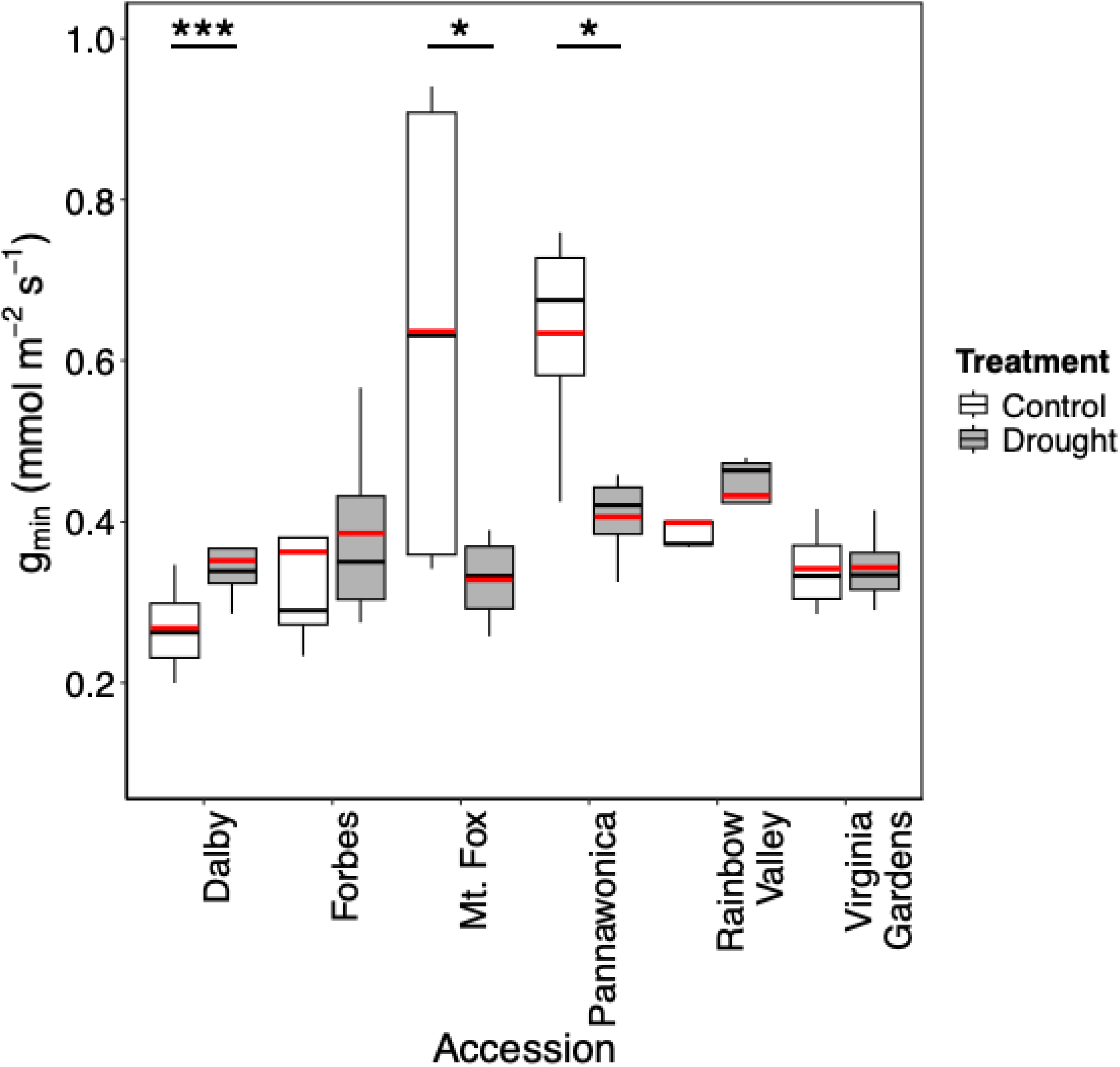
Box plots showing the variation in g_min_ across six accessions of *Themeda triandra* under control and drought treatments. Significant differences between treatments within accessions are denoted by asterisks (*p < 0.05, ***p < 0.001; paired t-test). The boxes represent the interquartile range (IQR), the black horizontal line within each box represents the median, and whiskers extend to 1.5 × IQR (n = 4). Red line inside the boxes represent mean.

For droughted plants g_min_ also varied among accessions (ANOVA, p = 0.041), albeit sufficiently weakly that no pairwise differences were statistically significant after post hoc correction (Tukey HSD, all p > 0.05). One comparison was very nearly statistically significant, with plants from Rainbow Valley tending to have higher g_min_ than those from Mt. Fox (Tukey HSD, p = 0.054).

Although there was no overall difference in g_min_ between control and droughted plants when pooled across all accessions, individual accessions differed in their g_min_ response to drought (accession × treatment interaction: ANOVA, p < 0.001), with two exhibiting pronounced reductions in g_min_ (Mt. Fox and Pannawonica; paired t-test, p = 0.0162 and p = 0.0104, respectively), three showing no significant difference in g_min_ with treatment, and one (Dalby) exhibiting increased g_min_ (paired t-test, p < 0.001). The asterisks in Figure 3 identify these accessions and the degree of significance of these shifts.

### Influence of climate-at-origin on g_min_ and its response to drought

To evaluate how climate-at-origin influences g_min_, we assessed relationships between g_min_ and both seasonal and annual climate variables. Our primary focus was on summer temperature and summer rainfall (Fig. 4), with relationships involving mean annual precipitation and mean annual temperature reported as supplementary results (Fig. S6).

**Figure 4.**
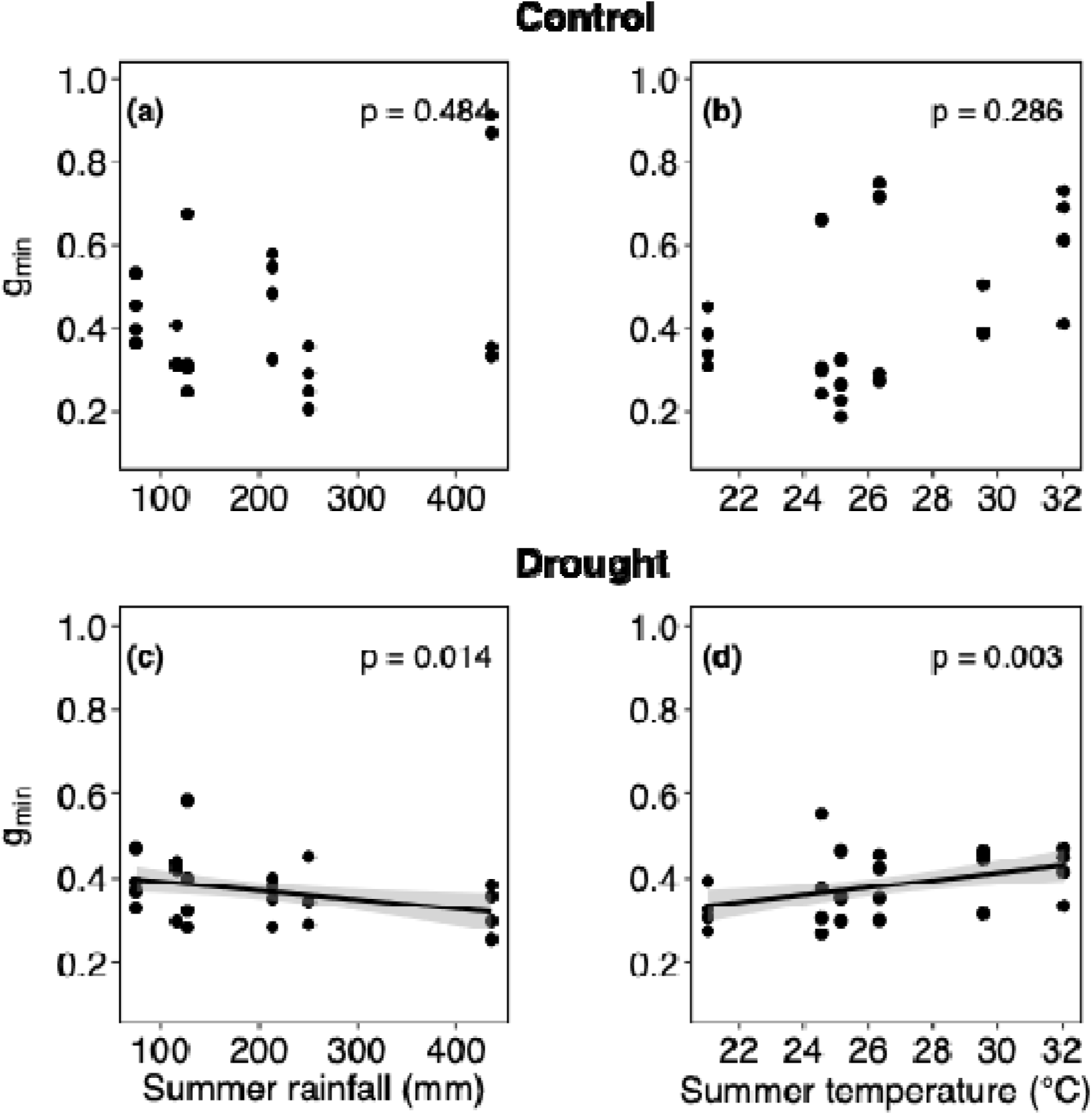
Relationship of g_min_ (mmol m^−2^ s^−1^) with climate-at-origin variables under control (a−b) and drought (c−d) conditions across *Themeda triandra* accessions. g_min_ is plotted against summer rainfall (a, c) and summer temperature (b, d). Data points represent individual measurements. Black lines and shaded areas depict fitted values and 95% confidence intervals from linear mixed effects models, with accession included as a random effect.

Under control conditions, g_min_ showed no relationship with either summer rainfall (p = 0.484; Fig. 4a) or summer temperature (p = 0.286; Fig. 4b). Similarly, g_min_ showed no relationship to either AP or AT (both p > 0.12; Figs S6a,b). By contrast, clear climate-linked patterns emerged under drought conditions, g_min_ being on average higher in plants from drier sites (Figs 4c, S5c; p = 0.014 and 0.002 respectively), such that a reduction in summer rainfall from 436 mm to 75 mm corresponded to a 16.5% increase in g_min_. On average, g_min_ of droughted plants was also higher in plants from sites with warmer summers (p = 0.003; Fig. 4d), such that an increase in summer temperature from 21 °C to 32 °C corresponded to a 23.5% increase in g_min_. There was no relationship between g_min_ and MAT (p = 0.961; Fig. S6d).

To assess plasticity in cuticle conductance, we quantified the percentage change in g_min_ from control to drought (%Δg_min_). Contrary to our expectation that accessions from hotter or drier environments would show greater reductions in g_min_, %Δg_min_ was not statistically related to any of summer rainfall, summer temperature, AP or AT (Figs. 5a, b and S6a, b; all p > 0.05).

**Figure 5.**
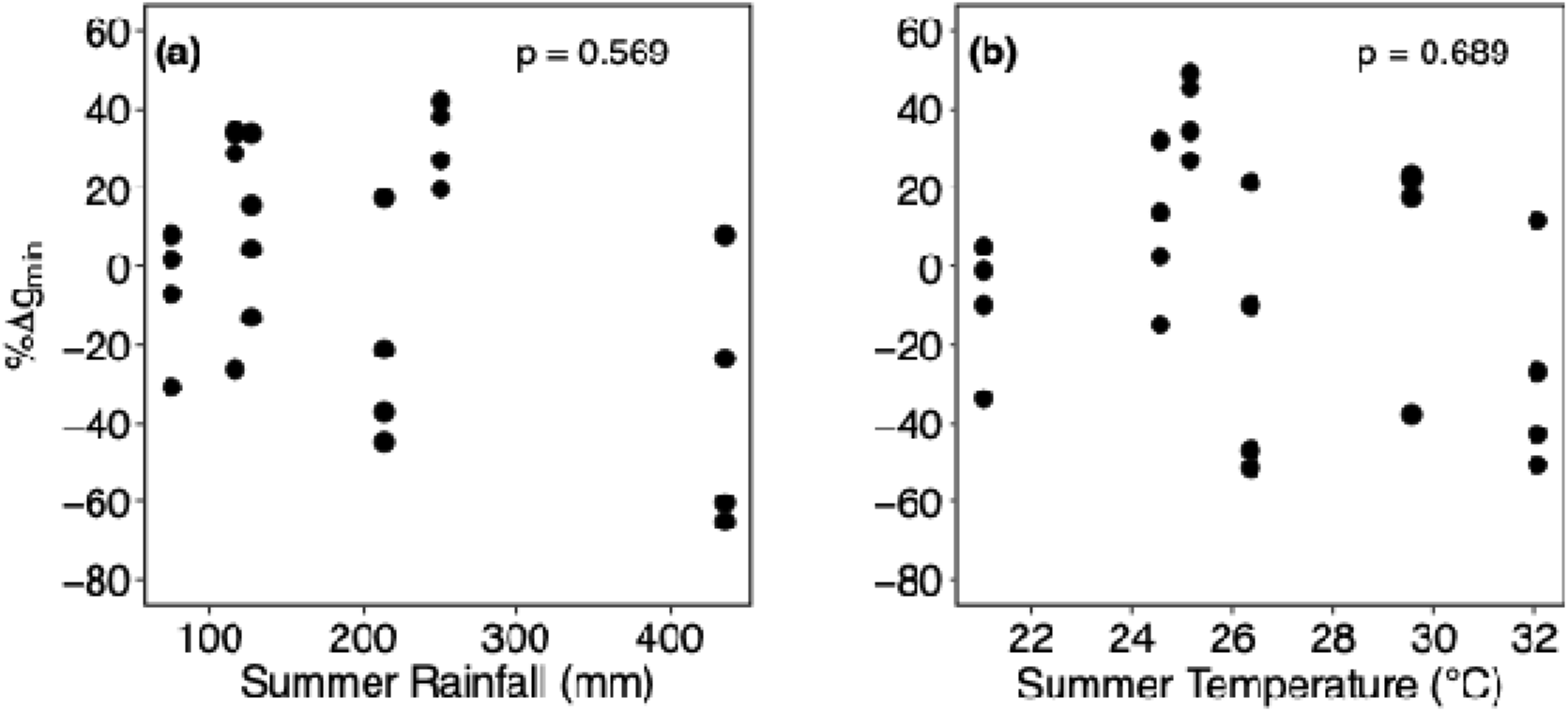
Relationship between %Δg_min_ (mmol m^−2^ s^−1^) and climate-at-origin variables across *Themeda triandra* accessions. (a) Summer rainfall (mm) and (b) summer temperature (°C) are shown as predictors. Each point represents an individual observation. Relationships were evaluated using linear mixed-effects models, with accession included as a random effect.

### Temperature dependence of g_min_ across accessions

The “textbook” response of g_min_ to temperature was not observed in our study (i.e., we did not observe a consistent pattern of near-constant g_min_, rapidly increasing at some threshold temperature, T_p_). Overall, the response of g_min_ to increasing temperature was broadly similar across accessions (Fig. 6, a−f). While most accessions (Mt. Fox, Pannawonica, Rainbow Valley and Virginia Gardens) showed a relatively flat response with slightly decreasing g_min_ as temperature increased, plants from Dalby and Forbes displayed a modest rise in g_min_ around 35−40 °C before eventually declining at higher temperatures, like the other accessions. A segmented regression model applied to individual replicates revealed no consistent evidence of a distinct breakpoint across accessions. This statistical outcome aligns with the visual inspection of the plots, where g_min_ appeared largely stable or declined gradually across the experimental temperature range.

**Figure 6.**
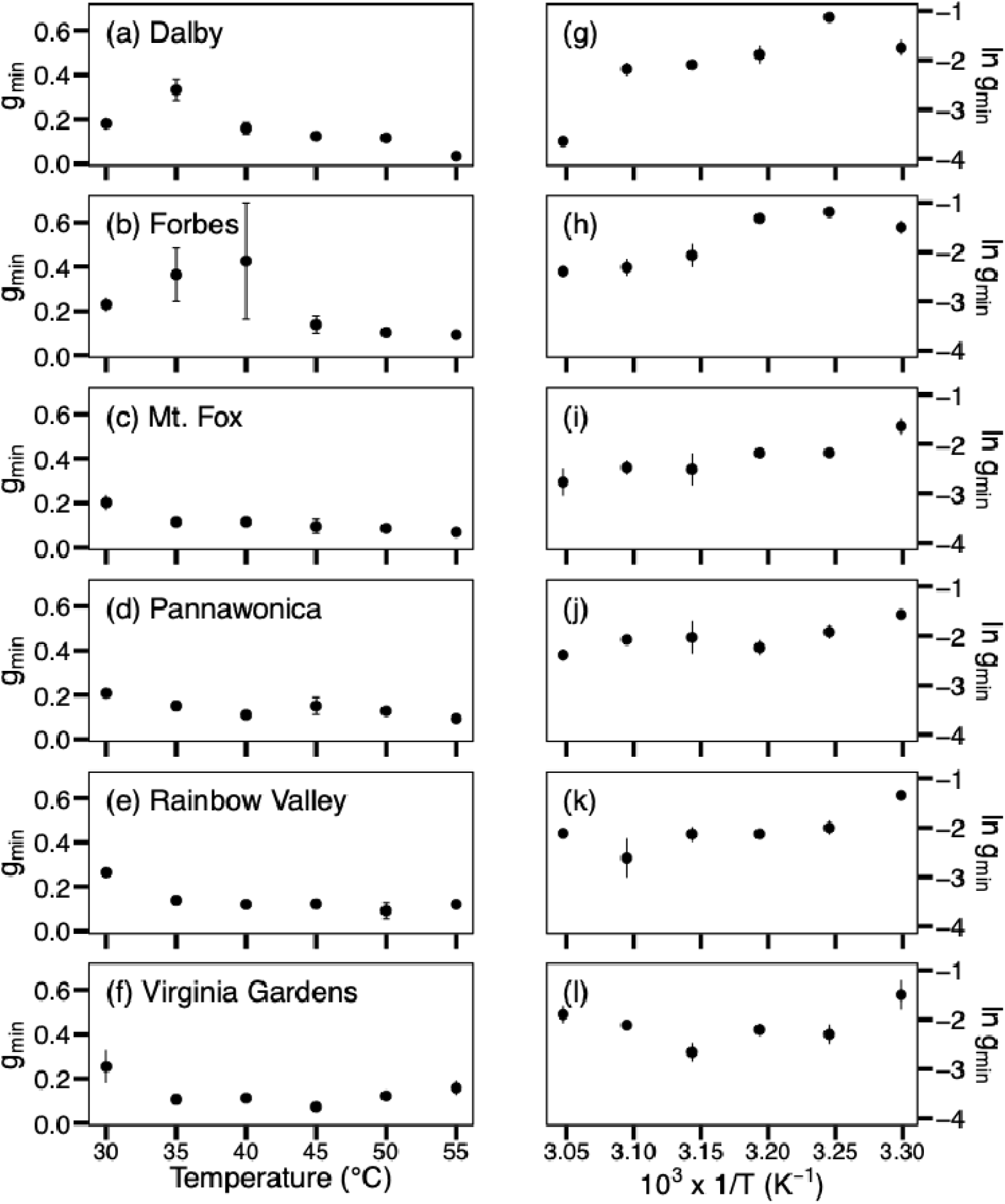
Temperature dependence of g_min_ (a−f) and corresponding Arrhenius plots (g−l) across a temperature range of 30−55□ at a constant absolute humidity of 18 g(H_2_O)m^−3^

## Discussion

### Intraspecific variation in g_min_ responses to drought

As the plants were well-watered before a single, short drought cycle, the treatment primarily reflects acute physiological plasticity rather than long-term structural acclimation. Brief water deficits can rapidly induce cuticular adjustments via abscisic acid (ABA) signalling and reorganisation of the wax biosynthetic pathways in plants, often within hours to days (Shepherd & Wynne Griffiths, 2006, Macková *et al*., 2013, Yang *et al*., 2020). Similar short term wax shifts have been documented in *Nicotiana glauca* (Cameron *et al*., 2006).

This study highlights the impact of short-term drought stress on g_min_ across six accessions of *T. triandra,* revealing substantial within-species variability in responses. Although the overall effect of drought on g_min_ was not statistically significant at the species level (Fig. 2), the direction and magnitude of responses differed substantially among accessions (Fig. 3), pointing to accession-specific strategies in cuticular water regulation. The lack of association between leaf water potentials and %Δg_min_ suggests that differences in cuticular conductance among accessions were not driven by unequal drought severity but instead reflect inherent variation in cuticle function and plasticity.

The significant reductions in g_min_ in response to drought observed in accessions from Mt. Fox (48%, p = 0.0162) and Pannawonica (36%, p = 0.0104) align closely with the classical drought-adaptive responses. While the drought response of g_min_ was not directly correlated with the accessions’ climate-at-origin, these two regions share ecological features—seasonally intense or prolonged drought, high evaporative demand, and unpredictable rainfall. These climate features may favour physiological strategies that reinforce the cuticle under water deficit. In these cases, the decrease in g_min_ may therefore reflect inducible cuticular responses, such as increased wax deposition or structural adjustments that strengthen the water barrier.

However, these patterns were not universal. The Dalby accession exhibited a 31% increase in g_min_ (p < 0.001) under drought stress—a response that diverges from the typical drought-induced reduction. This increase may reflect a plastic, rather than adaptive, response to drought—potentially involving alterations in cuticular wax composition that do not enhance water barrier function—as observed in grape berries under water deficit (Dimopoulos *et al*., 2020). Notably, Dalby had lower g_min_ than the other accessions under control conditions and, even with this increase, its absolute g_min_ remained relatively low. This suggests that plants from Dalby may possess inherently lower cuticular permeability overall, and that the observed increase may still fall within a functionally conservative range.

In contrast, in plants from Forbes, Rainbow Valley and Virginia Gardens, g_min_ did not differ between control and drought treatments. This suggests a high degree of drought tolerance or a mechanism of homeostasis that stabilizes physiological performance under fluctuating water availability. Such accessions might possess robust cuticular barriers that maintain consistent conductance levels regardless of external stressors. Alternatively, these accessions may rely on non-cuticular mechanisms—such as deep or extensive roots systems or osmotic adjustment—to cope with water stress. For instance, efficient root systems can enhance water uptake (Wasson *et al*., 2012), and osmotic adjustment can maintain cell turgor (Turner, 2018) and help plants tolerate water limitation (DaCosta & Huang, 2006). Osmotic adjustment involves the accumulation of solutes in plant cells (Hsiao *et al*., 1976), which aids in retaining water and sustaining physiological processes during periods of limited water availability. Further research in *Themeda* is needed to investigate these processes.

### Climatic correlations with g_min_ and %Δg_min_

*T. triandra* accessions revealed nuanced climate relationships with g_min_, underscoring the complexity of adaptive trait variation across environmental gradients. Under control conditions, g_min_ showed no association with either annual (AP, AT) or seasonal (summer temperature, summer rainfall) climate variables. This finding contrasts with our hypothesis that accessions from hotter and drier regions would exhibit inherently lower g_min_ to limit water loss. Instead, it suggests that cuticular permeability in this species under control conditions is not tightly linked to climate-at-origin, especially under control conditions. A comprehensive review by Duursma *et al*. (2019) found that g_min_ varies widely among species, but this variation is not clearly related to climate-of-origin. Very few studies have identified significant relationships between g_min_ and environmental variables. In a study on 42 conifer species (Brodribb *et al*., 2014) found a relationship between g_min_ and rainfall at the species’ origin, specifically for the driest quarter of the year, but their methodology involved gas exchange measurements rather than mass loss from detached leaves.

Under drought conditions, however, g_min_ displayed clear climate-linked trends, being typically higher in plants from drier and hotter sites (Figs 4c,d). This runs counter to classical expectations of drought adaptation, which predict reduced g_min_ as a water saving mechanism (Smith *et al*., 2006). Instead, it suggests that in hot, dry climates, maintaining moderate levels of cuticle permeability may serve alternative functions—such as leaf surface cooling via residual transpiration (Marchin *et al*., 2023). However, it is important to note that the magnitude of these trends was relatively modest—across the range of observed summer rainfall (75−436 mm), g_min_ increased by only 16.5%, and across the summer temperature range (21−32 °C), by 23.5%. Notably, MAT was not a significant predictor of g_min_ under either control or drought conditions, emphasizing that seasonal climate variables may be more informative than annual averages for understanding trait-environment relationships in this species.

The climate-linked pattern of g_min_ under drought—specifically, its positive relationship with summer rainfall in fig. 4c—may reflect an evolved alignment of trait means with long-term climate. However, within individual accessions, the short-term drought responses were not always consistent with this trend. For example, Pannawonica—an arid origin accession—showed a reduction in g_min_ under drought. In contrast, Dalby—from a wetter site—showed an increase in g_min_ under drought. This suggests that trait means (shaped by climate adaptation), and trait plasticity (plastic responses to acute stress) are not always aligned. Trait means likely reflect long-term selective pressures at population level, whereas plastic responses to short-term drought may be shaped by local adaptive strategies, trade-offs, or even stress-related responses. Recognizing that trait means and plasticity are distinct and potentially uncorrelated axes of variation is important for interpreting plant responses to environmental change. Future work should therefore aim to disentangle the genetic, physiological and ecological bases of both axes, ideally by pairing analyses of traits in relation to long-term climate with experiments designed to capture short-term plasticity under controlled-stress scenarios.

We also assessed the plasticity of g_min_ by examining the percentage change from control to drought %Δg_min_, under the expectation that accessions from harsher environments might show greater reductions to conserve water. However, %Δg_min_ was not related to any climatic variable (Fig. S7), suggesting that responsiveness of cuticular conductance to drought is not predictable by climate in this species. This could imply that g_min_ plasticity is under weak environmental selection, for example, or perhaps that traits—such as osmotic adjustment or rooting depth—buffer water loss during drought more effectively.

Taken together, these findings highlight that drought-expressed g_min_ is more strongly shaped by climate-at-origin than baseline g_min_, particularly in relation to summer climate extremes. Yet, the direction of this relationship is opposite to classical water conservation models, pointing to the multifaceted roles of cuticle conductance in plant adaptation.

### Temperature dependent behaviour of g_min_

The temperature response of g_min_ is often described as biphasic, monotonically increasing, where conductance rises sharply beyond a transition point—typically around 35−40 °C. However, not all species follow this pattern. For instance, *Phoenix dactylifera* shows an invariant relationship between g_min_ and temperature (Bueno *et al*., 2019). Additionally, in a broader survey of tropical trees, only seven out of twenty-four species displayed this biphasic response, while most showed weakly decreasing, U-shaped, or invariant trends (Slot *et al*., 2021). Our investigation into temperature response of g_min_ in *T. triandra* accessions revealed a general trend of gradual decline in g_min_ with increasing temperature across most accessions. Studies report that g_min_ can decrease, remain invariant, or show a U-shaped response with increasing temperature, depending on species and environmental context (Duursma *et al*., 2019, Slot *et al*., 2021, Garen & Michaletz, 2025). In some cases, acclimation to higher growth temperatures leads to a reduction in g_min_. For example, Zailaa *et al*. (2025) found that Quercus shows a reversible response to temperature, with g_min_ decreasing from 25−45 °C and then returning back to original steady state. Similarly, *T. triandra* shows a decreasing trend in g_min_, although we do not know if this response is a reversible one. While direct studies on *T. triandra* or grasses in general are lacking, there is broad evidence among other species showing that g_min_ can show variable temperature-dependent trends.

Two accessions (Dalby and Forbes) showed a modest increase in g_min_ at intermediate temperatures, followed by a subsequent decrease—a pattern not observed in other accessions. Such non-monotonic trends may arise from reversible changes in wax fluidity. This atypical pattern suggests the involvement of temperature-sensitive physiological pathways or distinct differences in cuticular lipid composition, which are known to play a crucial role in regulating water permeability (Schönherr *et al*., 1979).

Arrhenius plots of g_min_ did not reveal a clear biphasic pattern, indicating the absence of a distinct phase transition across accessions and suggesting that cuticle remains relatively stable across the temperature range tested. We used fresh cohorts from well-watered plants for each temperature treatment and leaves from the higher temperature runs (50 °C and 55 °C) did not appear extremely dry or noticeably different from those from lower temperatures. This suggests that markedly reduced water status was unlikely to have prevented an increase in g_min_, although relative water content (RWC) was not measured and subtle differences cannot be excluded. Such differences could potentially contribute to the absence of a clear T_p_. However, these findings imply that *T. triandra* may lack a distinct thermal threshold for cuticle conductance, and that any structural adjustments in the cuticle occur progressively or are buffered by other physiological or biochemical mechanisms. The absence of a common temperature response pattern across accessions further highlights the complexity of thermal strategies within the species.

## Conclusion

Our findings provide new insights into the drought and thermal responses of *T. triandra* leaf cuticles and their relationship with environmental conditions, highlighting potential mechanisms driving these patterns. The observed variability in g_min_ could inform the selection of resilient genotypes for ecological restoration or breeding programs aimed at mitigating the impacts of climate change. Future work should focus on the biochemical composition of cuticular waxes and their role in modulating g_min_ under stress conditions. Comparative transcriptomics or metabolomic analyses may also elucidate the genetic underpinnings of these diverse responses. Additionally, field studies assessing the interplay between cuticular properties, drought, and temperature in natural populations would provide valuable context for our controlled experiment findings. While this study provides important insights, the controlled conditions of the experiments of do not fully replicate the complexity of field environments. Further research is needed to validate these findings across broader spatial and temporal scales. Incorporating long-term monitoring of *T. triandra* populations under varying climatic conditions will help refine our understanding of the evolutionary and ecological significance of cuticular adaptations in response to environmental change.

As climate change drives more frequent droughts and heatwaves, understanding how different species/accessions regulate water loss through their cuticles and withstand high temperatures is key to strengthening ecosystem resilience. Additionally, examining how cuticular permeability interacts with other drought responses—such as osmotic regulation and root structure—could improve predictions of plant survival under future climatic conditions. To turn these insights into practical conservation strategies, future studies should expand to include other grassland species and assess their responses in long-term field trials.

## Supplementary information

Supplementary data consist of the following: Table S1: Climate and origin of the *Themeda triandra* accessions used in the study. Figure S1: Soil moisture across three days of drought stress. Figure S2: Midday leaf water potential (MPa) under control and drought. Figure S3: Relationship between midday leaf water potential and %Δg_min_. Figure S4: Photograph of DroughtBox. Figure S5: Performance of DroughBox in maintaining stable conditions. Figure S6: Relationship between g_min_ and annual climatic variables. Figure S7: Relationship between %Δg_min_ and annual climatic variables.

## Author contributions

AM, VJ and IJW designed and conducted the experiment, analysed the data, and drafted the initial manuscript. IJW, BC, VJ and CJB assisted with data interpretation and manuscript revisions. All authors reviewed and approved the final manuscript.

## Conflict of interest

The authors declare no conflict of interest.

## Funding

This research was supported by funding from the Australian Research Council (ARC) Centre of Excellence (CoE) for Plant Success in Nature and Agriculture (CE200100015). Anu Middha was supported by a Postgraduate Research Scholarship from Western Sydney University and a PhD top-up scholarship from the ARC CoE.

## Acknowledgements

This work was supported by funding from the Australian Research Council and from Hawkesbury Institute for the Environment, Western Sydney University (WSU). We are very grateful to Dr Craig Barton, WSU, for constructing the DroughtBox and providing training in its use. We also thank Erick Calderon-Morale for his help in using the DroughtBox during the early stages of this work.

## Data availability

The raw data supporting this study have been uploaded to Zenodo (doi: 10.5281/zenodo.16884594) in restricted mode and will be made publicly accessible upon acceptance.

## Notes

https://doi.org/10.5281/zenodo.16884594

### Competing Interest Statement

The authors have declared no competing interest.

### Summary of Updates

A new analysis was added to ensure all plants were drought stressed. The efficiency of Droughtbox in maintaining stable conditions was demonstrated too.

