## Supplementary Information for "Drought, thermal response and climate-patterning in cuticular conductance of the widespread C4 grass, *Themeda triandra*"

Table S1. Summary of geographic origin and historical climate parameters for the six *T. triandra* accessions included in this study. Climate data (1981-2010) are from the CHELSA (Climatologies at high resolution for the earth’s land surface areas) data of downscaled model output temperature and precipitation estimates of the ERA-Interim climatic reanalysis to a high resolution of 30 arc sec. AT, Annual Temperature (℃); AP, Annual Precipitation (mm)

| **Accession** | **State** | **Latitude** | **Longitude** | **Summer temperature (℃)** | **Summer rainfall (mm)** | **AT (℃)** | **AP (mm)** |
| --- | --- | --- | --- | --- | --- | --- | --- |
| Dalby | QLD | -27.1148 | 151.1817 | 25.1 | 250.4 | 19.6 | 627 |
| Forbes | NSW | -33.4076 | 147.9654 | 24.5 | 127.1 | 17 | 490 |
| Mt. Fox | QLD | -19.0013 | 145.4731 | 26.3 | 435.9 | 22.7 | 712 |
| Pannawonica | WA | -21.6449 | 116.3233 | 32.0 | 213.8 | 26.9 | 387 |
| Rainbow Valley | NT | -24.2376 | 133.5536 | 29.5 | 116.4 | 22.1 | 281 |
| Virginia Gardens | SA | -34.8538 | 138.6205 | 21.0 | 75.1 | 16.2 | 668 |

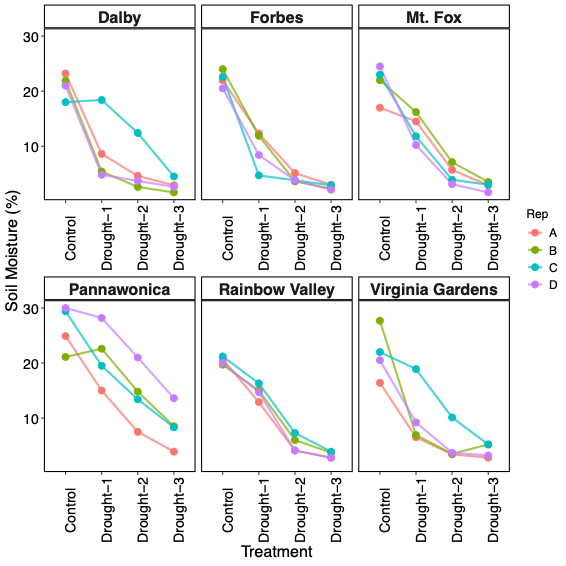

Figure S1. Changes in soil moisture (%) across three days of drought stress for six accessions of *T. triandra*. Each line represents an individual biological replicate showing consistent declines in soil moisture following drought imposition. Different replicates (A, B, C, D) of the accessions are shown by different colours.

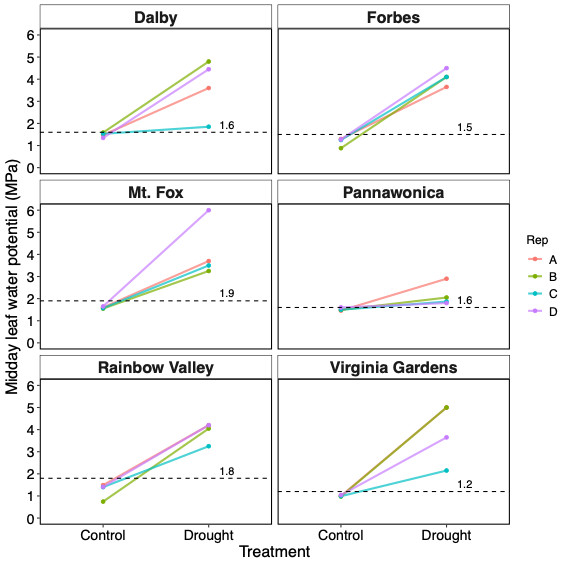

Figure S2. Midday leaf water potential (MPa) under control and day-3 drought conditions for six *T. triandra* accessions. Each line connects paired measurements from the same biological replicate. Different replicates (A, B, C, D) of the accessions are shown by different colours. The horizontal line in each panel represents the accession-specific water potential at turgor loss point, which was used as a threshold of drought stress individual plants needed to exceed to be included in the drought response analysis.

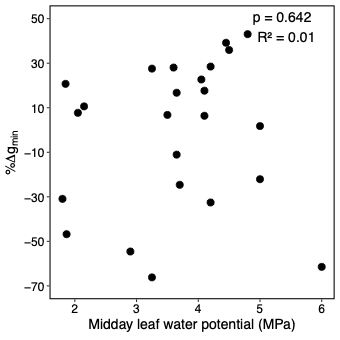

Figure S3. Relationship between midday leaf water potential and %∆g_min_ under drought across individual plants. Each point represents a single replicate. The absence of a significant relationship (p = 0.642, R^2^ = 0.01) indicates that variation in drought severity among individuals did not influence the plants’ response of g_min_ under drought.

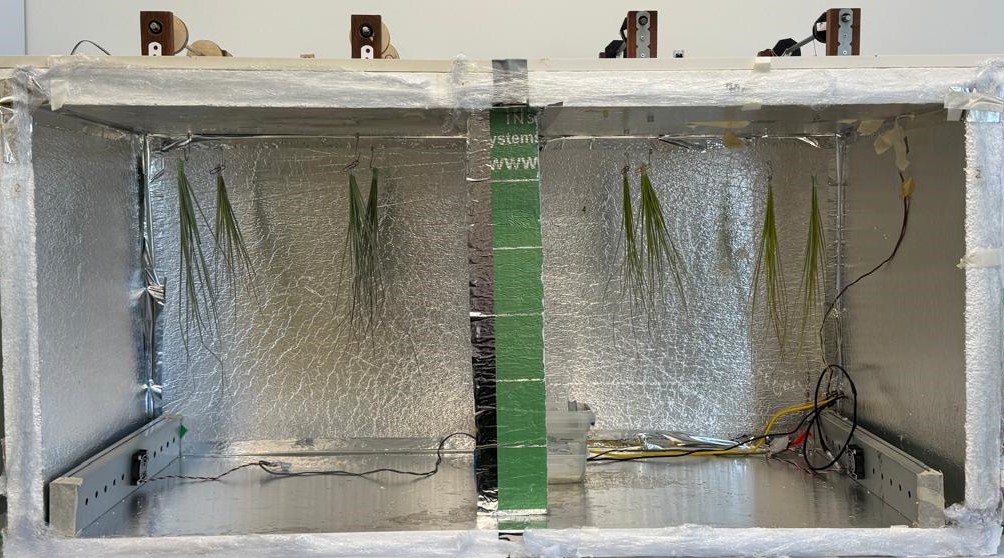

Figure S4. Photograph of the DroughtBox (front door removed) with samples set up and prepared to initiate a leaf-drying run.

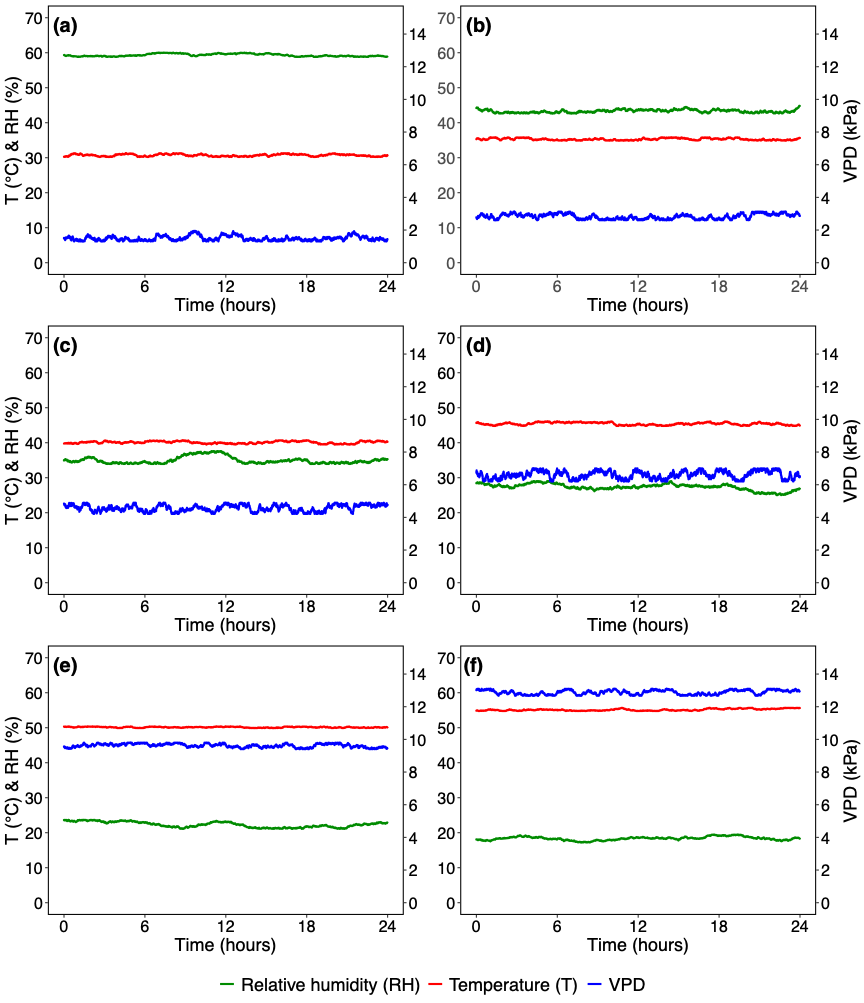

Figure S5. Performance of the DroughtBox in maintaining stable conditions across temperature treatments. Panels (a-f) show the regulation of temperature (T, red), relative humidity (RH, green) and vapour pressure deficit (VPD, blue) during 24-hour test runs at programmed temperatures of (a) 30 °C, (b) 35 °C, (c) 40 °C, (d) 45 °C, and (f) 55 °C. Corresponding increase in VPD ensured constant absolute humidity across different temperatures. The system maintained steady values of T, RH, and VPD throughout each test period, confirming its capacity for precise environmental control.

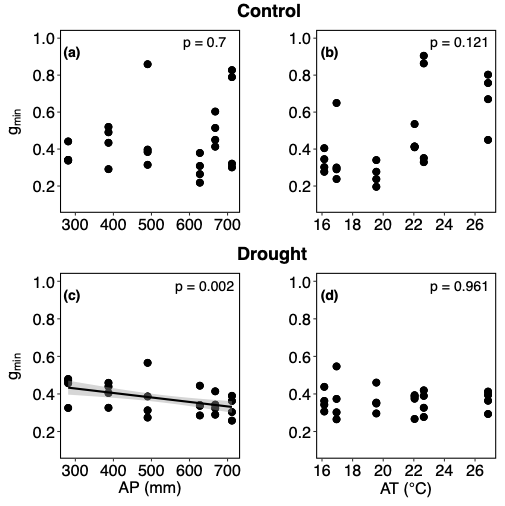

Figure S6. Relationships between g_min_ (mmol m^-2^ s^-1^) and annual climate variables across *T. triandra* accessions under control (a-b) and drought (c-d) conditions. Panels show g_min_ as a function of annual precipitation (AP) and annual temperature (AT). Each point represents an individual observation. Relationships were evaluated using linear mixed-effects models, with accession included as a random effect.

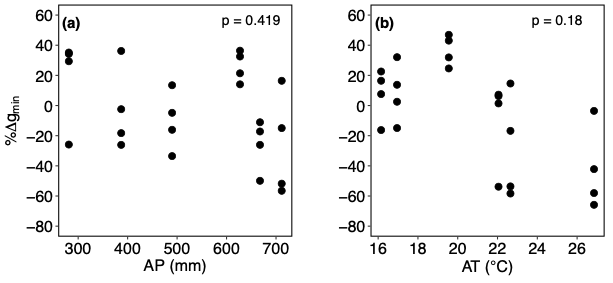

Figure S7. Relationship between %∆g_min_ (mmol m^-2^ s^-1^) and annual climate variables: (a) annual precipitation (AP) and (b) annual temperature (AT) across *Themeda triandra* accessions. Each point represents an individual observation. Relationships were evaluated using linear mixed-effects models, with accession included as a random effect.
